# Deep learning-based methods for individual recognition in small birds

**DOI:** 10.1101/862557

**Authors:** André C. Ferreira, Liliana R. Silva, Francesco Renna, Hanja B. Brandl, Julien P. Renoult, Damien R. Farine, Rita Covas, Claire Doutrelant

## Abstract

1. Individual identification is a crucial step to answer many questions in evolutionary biology and is mostly performed by marking animals with tags. Such methods are well established but often make data collection and analyses time consuming and consequently are not suited for collecting very large datasets.
2. Recent technological and analytical advances, such as deep learning, can help overcome these limitations by automatizing data collection and analysis. Currently one of the bottlenecks preventing the application of deep learning for individual identification is the need of hundreds to thousands of labelled pictures required for training convolutional neural networks (CNNs).
3. Here, we describe procedures that improve data collection and allow individual identification in captive and wild birds and we apply it to three small bird species, the sociable weaver *Philetairus socius*, the great tit *Parus major* and the zebra finch *Taeniopygia guttata*.
4. First, we present an automated method that allows the collection of large samples of individually labelled images. Second, we describe how to train a CNN to identify individuals. Third, we illustrate the general applicability of CNN for individual identification in animal studies by showing that the trained CNN can predict the identity of birds from images collected in contexts that differ from the ones originally used to train the CNNs. Fourth, we present a potential solution to solve the issues of new incoming individuals.
5. Overall our work demonstrates the feasibility of applying state-of-the-art deep learning tools for individual identification of birds, both in the lab and in the wild. These techniques are made possible by our approaches that allow efficient collection of training data. The ability to conduct individual identification of birds without requiring external markers that can be visually identified by human observers represents a major advance over current methods.

## INTRODUCTION

In recent years, artificial intelligence techniques, such as convolutional neural network (CNN), have caught the attention of ecologists. Such tools can automatize the analysis of various types of data, ranging from species abundance to behaviours, and from different sources such as pictures or audio recordings (reviewed in Christin, Hervet & Lecomte, 2019). CNNs are a class of deep neural networks that, contrary to other types of artificial intelligence methods that require hand-crafted feature extraction, automatically learn from the data the features that are optimal for solving a given classification problem (see Angermueller, Pärnamaa, Parts & Stegle, 2016; Christin et al., 2019; Jordan & Mitchell, 2015; LeCun, Bengio & Hinton, 2015 for a detailed introduction on deep learning). CNNs are thus particularly useful when many features for classification are needed.

In ecology, deep learning has been successfully and predominantly applied to identifying and counting animal or plant species from pictures. For example, Norouzzadeh et al. (2018) used a long term database of more than 3 million labelled pictures to train a CNN to automatically recognize 48 African animal species. This CNN can replace the need of human manual identification in future studies, thus promoting a more efficient data analysis. This and other examples (e.g. Rzanny, Seeland, Wäldchen & Mäder, 2017; Tabak et al., 2019) highlight the potentialities of deep learning for reducing human effort and increasing identification performance. Beyond species recognition, another promising application of CNNs is individual identification, which is crucial to many studies in ecology, behaviour and conservation (Clutton-Brock & Sheldon, 2010). Individual identification using deep learning has been the subject of extensive research in humans (e.g. Ranjan et al., 2018), and recently a handful of studies have applied it to other animal species (e.g. primates, Deb et al., 2018; pigs, Hansen et al., 2018; elephants, Körschens, Barz & Denzler, 2018). However, the application of deep learning to smaller taxa, and specifically birds, remains unexplored.

In birds, manual examination of pictures or video recordings of visually marked populations (e.g. using colour rings), are well established methods. However, relying on humans for individual identification and data collection is time consuming (Weinstein, 2018). In many cases the use of recently developed animal-tracking devices (e.g. GPS) and sensor technologies (e.g. RFID) can be used (reviewed in Krause et al., 2013). Yet, animal-borne tracking devices are also often limited when visual information on contexts and behaviours are important. For example, studying parental care in birds requires video recordings to visually identify which birds are providing care to the chicks and how often they do it, as well as to identify several other relevant behaviours and attributes, such as the type of food that parents are bringing to the chicks or distinguishing the purpose of the visit (e.g. to feed the chicks or to engage in nest maintenance activities). Thus, a major advance over current methods would be to automatically identify individuals while keeping the versatility of pictures and video recordings for behavioural data collection (which should in turn be automatized as well).

Several methods for automatic individual identification and other data extraction from pictures and videos of animals have been developed previously. For instance, Pérez-Escudero, Vicente-Page, Hinz, Arganda & de Polavieja (2014) proposed a multi-tracking algorithm capable of following unmarked fish in captivity from video recordings (which was later improved using deep learning; Romero-Ferrero, Bergomi, Hinz, Heras, & de Polavieja, 2019), whereas other computer vision-based methods that require tags or marks to assist with computer tracking and identification have been developed and applied in behavioural captivity studies (e.g. Alarcón‐Nieto et al., 2018). However, these methods are mostly limited to animals in captivity, either because they require standardized recording conditions (e.g. consistent background light, known number of individuals present in the recording) or the marks needed to assist identification are attached through gluing or through backpacks that are not suitable to be fitted to many animals, especially in the wild. Deep learning has the potential to overcome some of the limitations of the current automated methods, as it can identify individuals by relying only on their natural variance in appearance and be tolerant to spurious variation in the recording conditions.

A major challenge for the application of individual recognition using deep learning methods is the need of collecting extensive training data. Acquiring training data typically involves labelling images with the location and/or identity (or an attribute) of each individual. The amount of data required to train a CNN is expected to be proportionally dependent on the difficulty of the classification challenge, i.e. a bear and a bird would be easier to differentiate than two bears of the same species. Usually CNNs that achieve large generalization capability are trained over thousands to millions of pictures (Marcus, 2018). Such large datasets are required as usually CNNs have to generalize from the specific data that they have been exposed to during training. For example, if a CNN was trained to distinguish two bears of the same species with only pictures of the individuals lying down, difficulties may arise to identify those same individuals from new pictures taken when the animals were standing up. Additionally, if the pictures used for training were taken during a short period of time, it might lead the CNN to rely on superficial and temporary features for identification. For example, if pictures for training were taken when one of the individuals had a large wound or was going through moulting or shedding, it might result in a CNN that relies on those salient and temporary features and perform badly when predicting the identity of the individuals a few days later. Therefore, effectively making use of deep learning for individual identification, especially in the wild, requires an adaptive framework for collecting training data.

When working in captivity settings, such large labelled image datasets can be easily collected by temporarily and routinely isolating the animals in enclosures separated from the rest of the group while filming or photographing them. However for researchers working on wild populations collecting training data can become challenging and it might not be feasible to rely on traditional methods of individual identification for labelling the pictures. For example, in birds, relying on human observers and colour rings, to photograph and manually label enough pictures to implement CNN for individual identification, can become extremely costly and time-consuming. Furthermore, in longer-term studies, animals can change their appearance over time (e.g. changing from juvenile to adult plumage in birds) or new individuals may join the population (e.g. immigrants or recruited offspring). These cases require that the process of identifying individuals and labelling photos is routinely repeated. Therefore, relying on human observers for collecting labelled data in this type of systems might hinder the implementation of deep learning techniques for individual identification, or restrict its application to short-term projects.

Here, we provide guidance on how training data can be efficiently collected, both in captivity and in the wild, and on the subsequent steps required to train a CNN for individual identification. We demonstrate the feasibility of our approaches using data from two wild pit-tagged populations of birds from two different species, the sociable weaver *Philetairus socius* and the great tit *Parus major*, and a population of captive zebra finches *Taeniopygia guttata*.

We start by 1) focusing on the problem of efficiently collecting large training datasets. We provide simple and automated methods for collecting a very large number of labelled pictures by using RFID tags associated to camera traps (in the wild sociable weaver and the great tit populations) or by temporarily isolating the target individuals (in captive zebra finches). In all cases, we used low-cost RFIDs and low-cost cameras that can be programed to take labelled pictures of the birds’ back feathers. 2) We provide details of the data pre-processing and the training of an adequate CNN. 3) Subsequently, we evaluate the generalization performance of our CNNs to other circumstances by evaluating the ability of our models to predict the identity of the birds in pictures collected with different cameras and in contexts that differ from the ones used for collecting the training datasets. 4) Finally, we present a very simple approach to address the problems arising from the arrival of new and unmarked individuals to the population.

## METHODS

### Study populations

We collected pictures from a population of sociable weavers at Benfontein Nature Reserve in Kimberley, South Africa. For the great tits, pictures were collected from a population in Möggingen, southern Germany. For both species, birds were fitted with pit-tags as nestlings, or when trapped in mist-nets as adults and are habituated to artificial feeders that are fitted with RFID antennas, as part of two independent on-going studies in these populations. For the zebra finches, pictures were collected from a captive population housed in Möggingen, southern Germany. Birds were being kept in indoor cages in pairs and small flocks.

### Collecting training data

#### Sociable weavers

The collection of labelled pictures was automated by combining RFID technology (Priority1Design, Australia) with single-board computers (Raspberry Pi), cameras and artificial feeders. We fitted RFID antenna to small perches placed in front of plastic feeders filled with mixed seeds (Fig. 1a). Each RFID data logger was connected to a Raspberry Pi (detailed explanation of the developed setup is available at github.com/AndreCFerreira/Bird_individualID) which was connected to a Pi camera (we used Pi camera V1 5mp and V2 8mp). We programmed the Raspberry Pi to take a picture every time that a bird was detected on the RFID logger, with a 2 seconds gap between pictures. This interval was introduced in order to avoid having near-identical frames of the same bird that would increase overfitting of the CNN and jeopardize the generalization capability of the models (see “Convolutional neural networks” section). The Raspberry Pi was programmed to take pictures with different shutter speeds to account for variation in light conditions over the day. Each picture file was automatically labelled with the bird identity, known from the RFID logger and the time of shooting in the filename. Training data collection is therefore automatized by automatically linking the identity of the bird perching on the antenna while feeding to its pictures, without the need of human manual identification and annotation.

**Figure 1.**
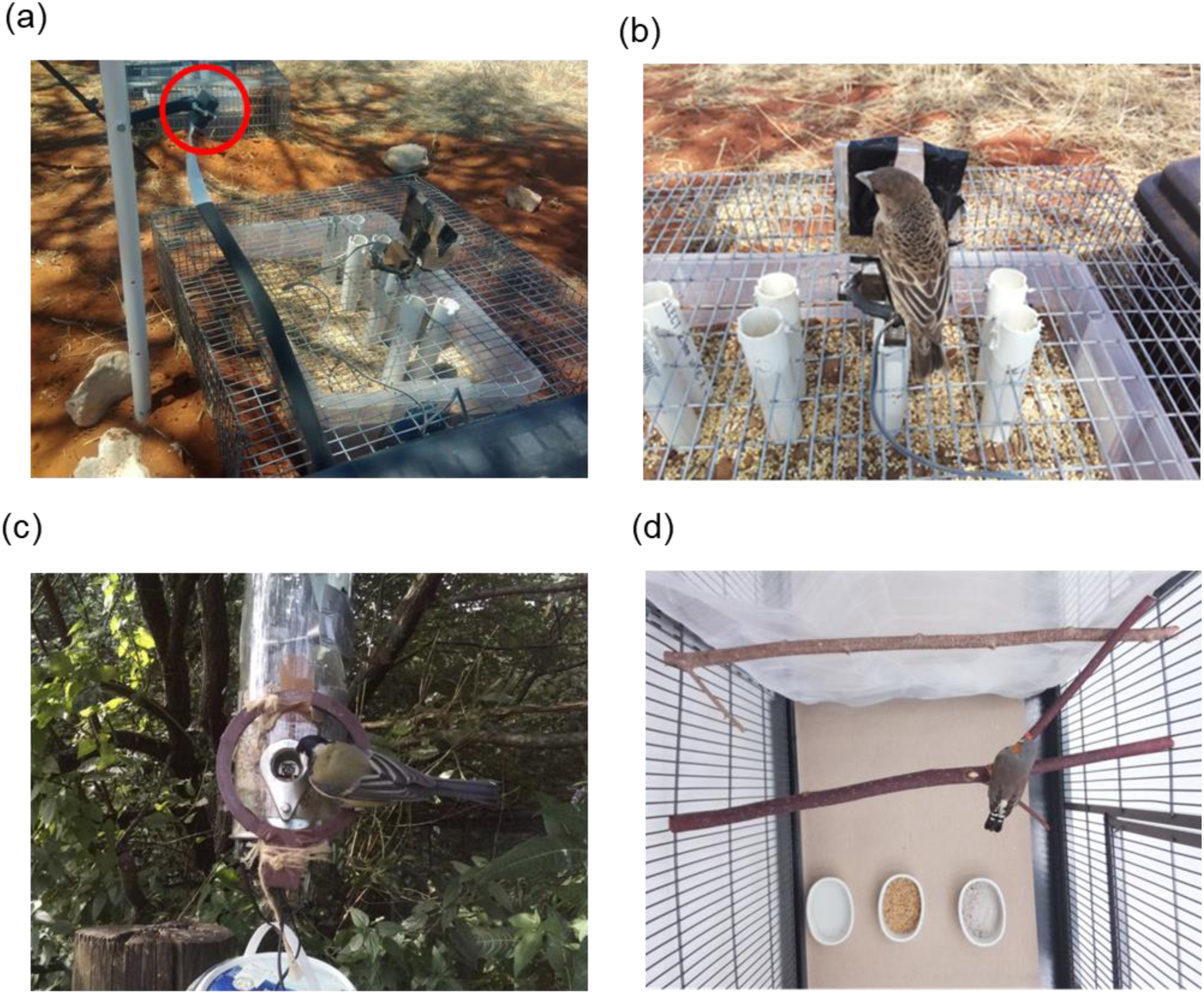
a) Pi camera (circled in red) positioned to record the back of the birds and b) the respective picture taken of a sociable weaver feeding while perched on the RFID antenna. c) Picture taken by the Pi camera of a great tit perching at the RFID antenna on a feeder and d) of a male zebra finch taken from inside the cage.

Three PI cameras and three feeders which were ca. two meters apart from each other were used. The cameras were positioned to take a picture from top perspective to enable to photograph both the scaled pattern of the back and wing feathers (Fig. 1b). The birds’ back was chosen as the distinctive mark since it is the body part that is most easily observed and recorded in multiple contexts (e.g. when perching at the feeders or building at the nest), making it a very versatile mark for applying an image classification algorithm in other contexts. Pictures were collected for 15 days during November and December 2018.

#### Great tits

We collected pictures of the individuals using a similar setup to the one described above, by placing a RFID antenna at an artificial feeder hanging on a tree branch (Fig. 1c). We used one single Pi camera and one feeder to collect pictures during seven days over the course of the last two weeks of August 2019.

#### Zebra finches

We temporarily divided aviaries into equally-sized partitions with a net to take pictures from individual birds without completely socially isolating them. We collected data from 10 zebra finches (five males and five females). In each partition, we placed two Raspberry Pi cameras to photograph (every two seconds) the birds sitting on the wooden perches (Fig. 1d). Each bird was recorded for four hours. Since we know which Raspberry Pi photographed which bird, we avoided the need to manually link the identity of the birds to the pictures.

#### Data pre-processing

To efficiently train a CNN, the regions in the pictures corresponding to the birds should be extracted from the background (second step of Fig. 2). Mask R-CNN (He, Gkioxari, Dollár & Girshick, 2017) was used to automatically localize and crop the bird out. For the sociable weavers, we used a Mask R-CNN model that has been trained on Microsoft COCO (Lin et al., 2014), a generalist dataset which includes pictures of birds and therefore is able to localize the sociable weavers in the pictures (see github.com/AndreCFerreira/Bird_individualID for details). For the great tits and zebra finches this Mask R-CNN model performed poorly and thus the model was re-trained by adding a new category (zebra finch or great tit, a different model for each species) and using pictures in which the region corresponding to the bird was manually delimited using “VGG Image Annotator” software (Dutta & Zisserman, 2019). Since manually labelling the regions of interest is time consuming, we started by training the model for 10 epochs with 200 pictures. If the model was found to perform badly, additional pictures were manually labelled and added it to the training dataset. This process was repeated until a satisfactory performance was achieved. For the great tits 500 pictures were used for training and 125 for validation (see “Convolutional neural networks” section below for explanation on training and validation datasets), for the zebra finch we used 400 pictures for training and 100 for validation.

**Figure 2.**
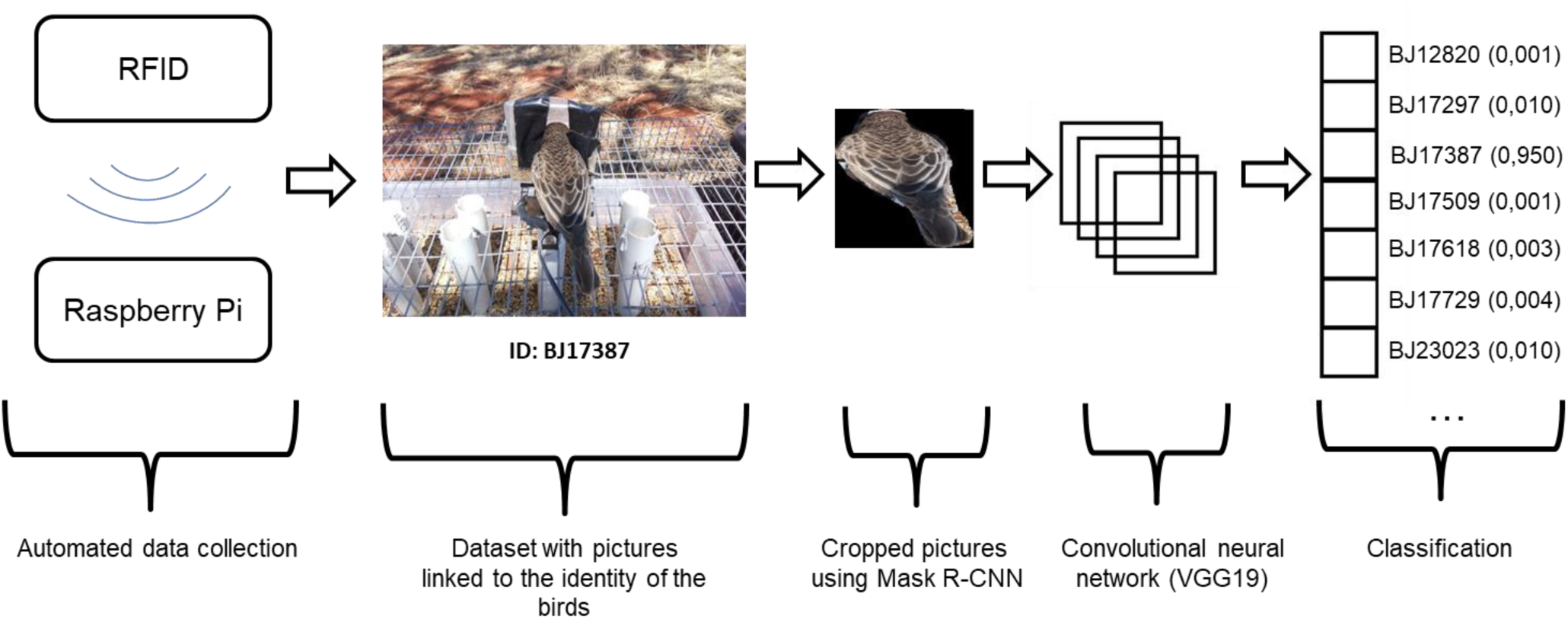
Sequential steps required for collecting data and training a convolutional neural network for individual identification.

From a total of 35 weavers detected at the RFIDs antennas we had 30 individuals with more than 350 pictures each that were used to train the classifier. In the great tit population, 77 birds were photographed including 10 with more than 350 pictures. These individuals were used to train a CNN. The remaining five weavers and 67 great tits (with less than 350 pictures) were used to address the issue of working in open areas where new individuals can constantly be recruited to the study population (see section “New birds” below). For the zebra finches we used all 10 individuals as our setup resulted in more than 2000 pictures for each bird.

### Convolutional neural networks

#### Sociable weavers

For 21 of the 30 selected sociable weavers, more than 1000 pictures were available and therefore we aimed at using 900 pictures for the training dataset and 100 pictures for validation dataset. For the 9 sociable weavers, for which we did not have 1000 pictures, we further avoided an imbalance in the dataset by first selecting 100 pictures for the validation dataset and then duplicating (through oversampling) the remaining pictures until 900 pictures were available for the training dataset (Buda, Maki & Mazurowski, 2018). We used 27038 unique pictures, 901.27±172.96 (mean±SD) per bird. The training dataset is the set of samples that the neural network repeatedly uses to learn how to classify the input images into different classes (in our case, different individuals). The validation dataset is used to compute the accuracy and loss (estimation of the error during training) of the model. This validation dataset is used to assess the learning progress of the neural network. As the network never trains or sees the validation data, this validation dataset can indicate if the model is overfitting the training data, i.e. if the model is “memorizing” the pictures instead of learning features that are key for recognizing the individuals.

To limit overfitting caused by having very similar pictures in the training and validation datasets, we used images for training and validation that were taken on different days. All pictures were normalized by dividing the arrays by 255 (0 to 1 normalization).

We used the VGG19 convolutional neural network architecture (Simonyan & Zisserman, 2014) and the weights of a network pre-trained on the ImageNet dataset (a dataset with more than 14 million pictures and 20000 classes, Deng et al., 2009). The main idea behind using networks pre-trained on other datasets is that features (such as colour or texture) that are important to distinguish multiple objects could also be useful to distinguish between individuals. The fully connected part of the VGG19 CNN network (i.e. the classifier part) were replaced by layers with random weights that fits our particular task of interest and the corresponding number of classes (30 individuals).

To further increase our training sample, we used data augmentation, which consists of artificially increasing the sample size by applying transformations to our existing sample. Using the data generator available in Keras, images were randomly rotated (from 0 to 40°) and zoomed (zoom range of 0.2). One 0.5 dropout layer was added just before the first dense layer to limit overfitting (see github.com/AndreCFerreira/Bird_individualID for details on the network architecture). We used a softmax activation function for the classifier. ADAM optimizer (Kingma & Ba 2014) was used with a learning rate of 1e-5. A batch size of eight was used since it has been shown that small batch sizes improve models’ generalization capability (Masters & Luschi, 2018). If there was no decrease in loss for more than 10 consecutive epochs we stopped training, and then retrained the model that achieved the lowest loss with a SGD optimizer and a learning rate 10 times smaller until there was no further decrease in the loss for more than 10 consecutive epochs. All analyses were conducted with python 3.7 using keras tensorflow 1.9, and on nvdia rtx 2070 gpu.

In an exploratory approach, and even though our model achieved ca. 90% accuracy with the validation dataset, the accuracy was significantly lower when generalizing to other contexts (see results). We suspected that such differences could be due to the lower quality of pictures collected in those other contexts (with different cameras, capture distances and conditions; see “Testing models” section). To account for this possibility we trained a model using the same setting parameters that yielded the best results, but applying Gaussian blur, motion blur, Gaussian noise and resizing transformations and a random combination of two of these four transformations (see github.com/AndreCFerreira/Bird_individualID for details on the transformations applied to the images) to each of the pictures of the dataset used to train the models in order to simulate the lower quality of the pictures taken in other contexts (Fig. 3). The idea is that even if the overall quality of the pictures in the dataset used for training slightly differs from pictures which are of interest for a research question, this training dataset can be transformed in order to be more similar to the pictures collected in distinct contexts for which the classifier could be applied on.

**Figure 3.**
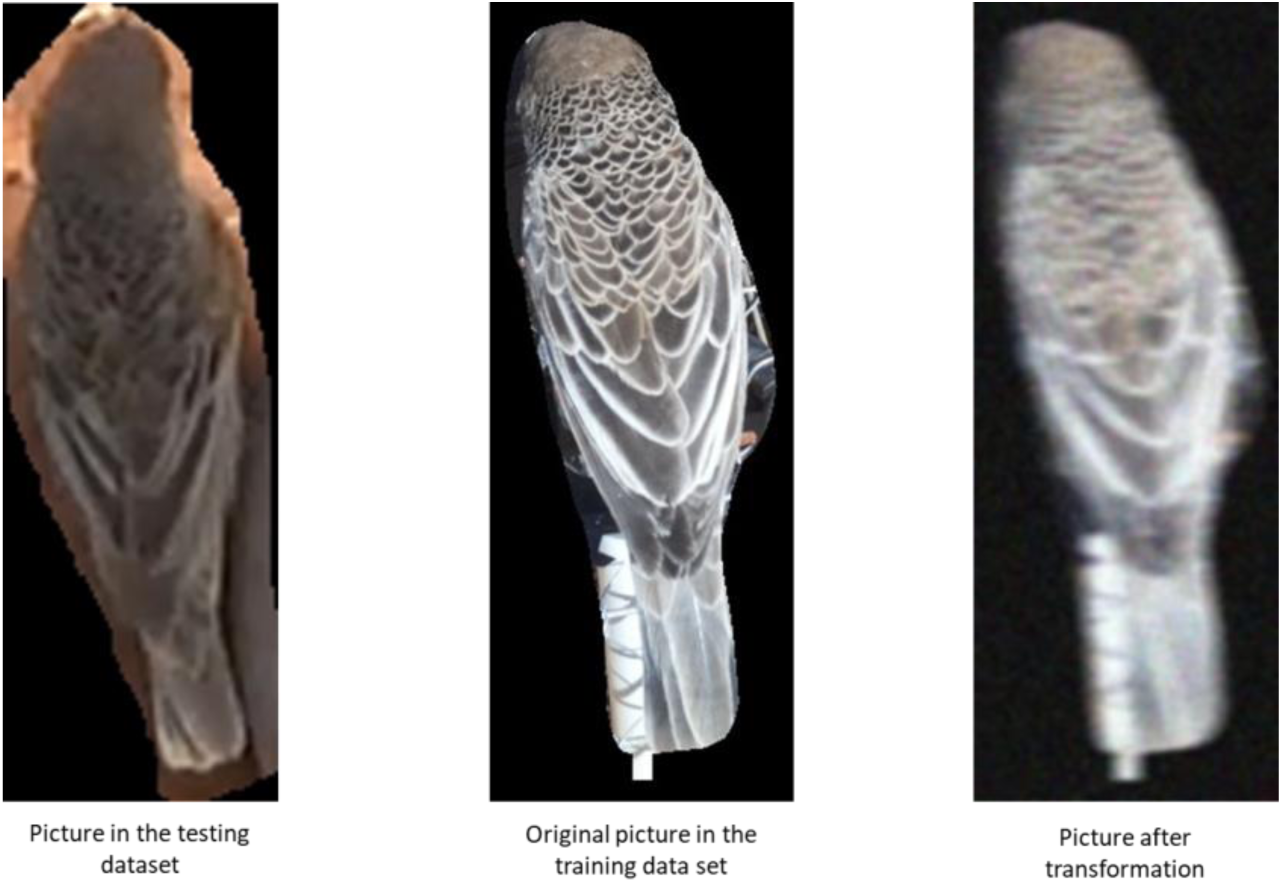
Comparison of the pictures’ quality in the testing dataset (on the left, see “Testing models” section below) with the training dataset (middle). On the right the same training pictures after applying a transformation to simulate the low-quality of the testing dataset.

#### Great tits

For the great tits we trained the CNN with 1000 pictures per bird, 900 pictures for training and 100 for validation. For birds with less than 1000 pictures (six birds) we did oversampling by creating copies of the pictures available following the same procedures as for the sociable weaver. We used 7605 unique pictures, 760.50±222.56 (mean±SD) per bird. Pictures in the validation dataset were also taken in different days from the pictures used for training.

The same architecture and hyperparameters as for sociable weavers were used, except that the dropout value was reduced to 0.2 as the model did not improve the accuracy from a random guess for 10 epochs when the dropout was at an initial value of 0.5. In addition to the zooming and rotation data transformations, horizontal and vertical flips were also used as the great tits, contrary to the sociable weavers, could be photographed from any orientation (as they perched all around the RFID antenna). Blur and noise transformations were not used as there were no differences in the overall quality of the pictures used for training and for testing the model generalization capability (see “Testing models” section).

#### Zebra finches

There were more pictures available per bird for the zebra finch than for the other species. However, the problem of collecting pictures in animals that are in confined enclosures is that a significant number of pictures could potentially be near-identical if the individuals stay motionless for long periods of time. In our case, all birds were generally active and visited all the places in their cage (i.e. all wooden perches, floor, water and food plates). Nevertheless, to avoid potential overestimation of the model’s accuracy, we used the pictures collected when the birds were in the left side of the cage for training and the pictures taken when the birds were on the right side of the cage for validation. Additionally, to create a diverse set of validation pictures, structural-similarity index measure (SSIM) (Wang, Bovik, Sheikh & Simoncelli, 2004) was used to make pairwise similarity comparisons between pictures. We started by randomly selecting a picture to include in the validation dataset. Additional pictures were then randomly sampled and used to compute the SSIM between the new picture and the ones already in the validation dataset and if the value was smaller than a threshold, these new pictures were included in the validation dataset. This process was repeated by sequentially comparing a new picture to all the ones already in the validation dataset until we reached 160 pictures per bird. The threshold value used (0.55) was empirically determined by trying different values and looking at the resulting datasets. For the training dataset, 1600 pictures of each bird were randomly selected without filtering for near-identical pictures. All birds had at least 1600, except for one that had 1197 for which oversampling was used by creating duplicates of randomly sampled 403 pictures.

Finally, the CNN was trained using the same procedures as for the great tits except that the dropout layer was set to 0.5 rather than 0.2.

### Testing models

#### Sociable weavers

To test the efficiency of our models, we collected images of the sociable weavers in different viewing perspectives, using different cameras and different contexts than the original feeding station setup. The aim was to evaluate the ability of our trained CNN to identify individuals in different experiments and contexts.

We used four different setups for testing. We filmed birds feeding in the same plastic RFID feeders but recorded using a Sony handycam (rather than Raspberry Pi camera), from two different perspectives: 1) close (95 pictures from 26 birds 3.65 ± 0.68 (mean ± SD; Fig. 4a) and 2) and far (71 pictures from 21 birds 3.43 ± 0.58; Fig. 4b). In addition, a plastic round feeder with seeds was positioned on the floor to record both from 3) a ground perspective (90 pictures from 28 birds 3.21 ± 1.21; Fig. 4c) and 4) a top perspective (83 pictures from 25 birds 3.32 ± 1.01; Fig. 4d).

**Figure 4.**
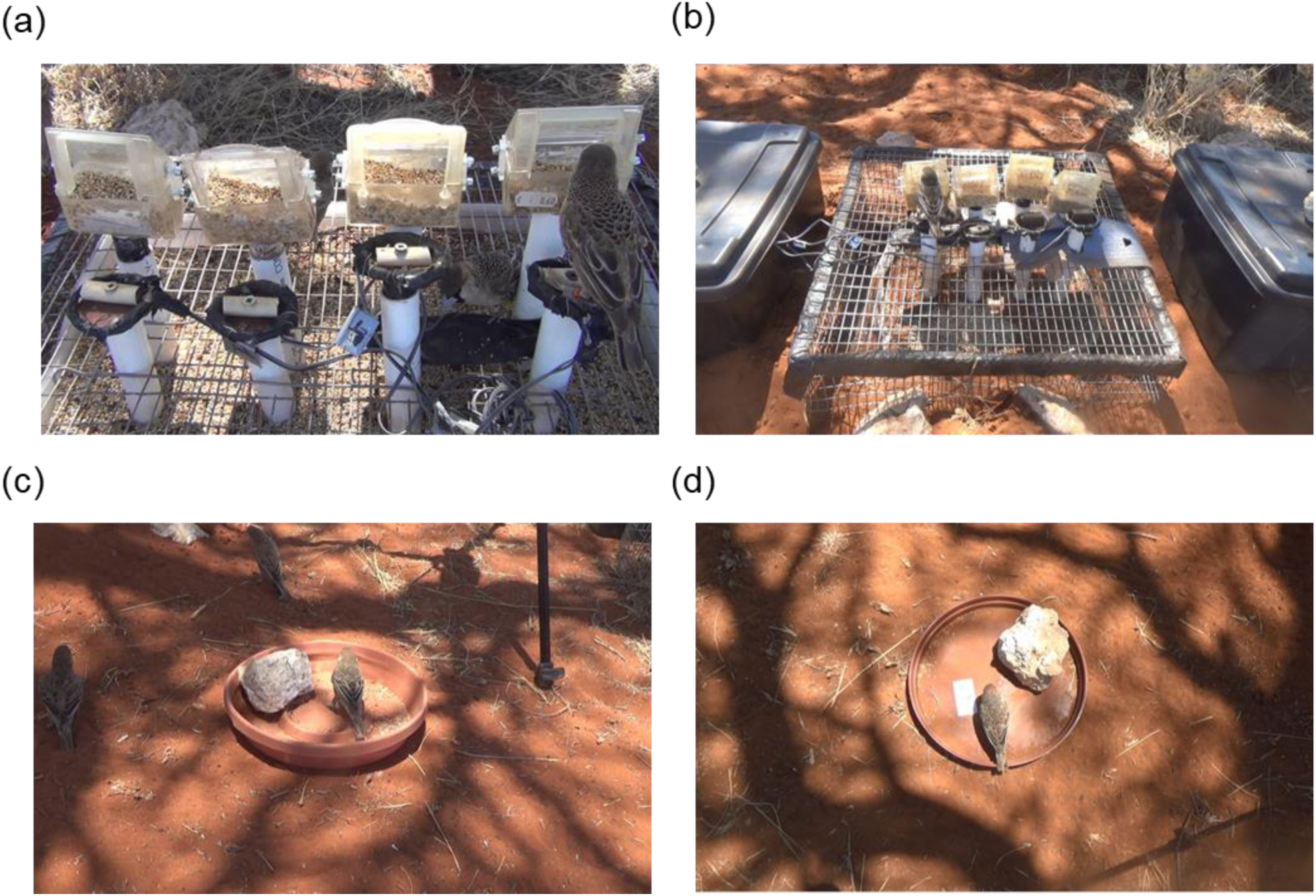
Example of pictures from the four different conditions used for the testing that were recorded at the feeders from the RIFD feeder setup from: a) close or b) far perspective, or directed at a feeding plate on the floor recorded from c) a ground perspective and d) a top perspective.

The birds were manually cropped out from pictures using imageJ (Schneider, Rasband & Eliceiri, 2012) and individually identified using their colour rings. The colour rings were then erased directly from the image to guarantee that the model did not use them for identification. Videos were recorded within the same time window as the training pictures collection and we aimed at extracting five non-identical frames per bird in which the back was fully visible, however this was not always possible for all birds as not all of them were recorded in these testing videos, or were not recorded long enough.

#### Great tits

We recorded birds feeding in a table from a top perspective with a Raspberry Pi camera (Fig. 5). Since these birds had no colour ring or any mark for visual identification, we identified them using their pit-tags by placing seeds on top of a RFID antenna in order to induce the birds to activate the RIFD antenna and obtain the identity of the birds feeding (similar to the pictures collected for training described above). Birds were recorded feeding on the table for 3 days but 4 out of the 10 birds in the training dataset did not use this new feeding spot. Additionally the number of pictures collected at this setup varied greatly between birds (from 2 to 38 pictures, mean: 15.7±11.3SD). We did not attempt to make a balanced dataset and, therefore, used all the 94 pictures collected at this new feeding set-up.

**Figure 5.**
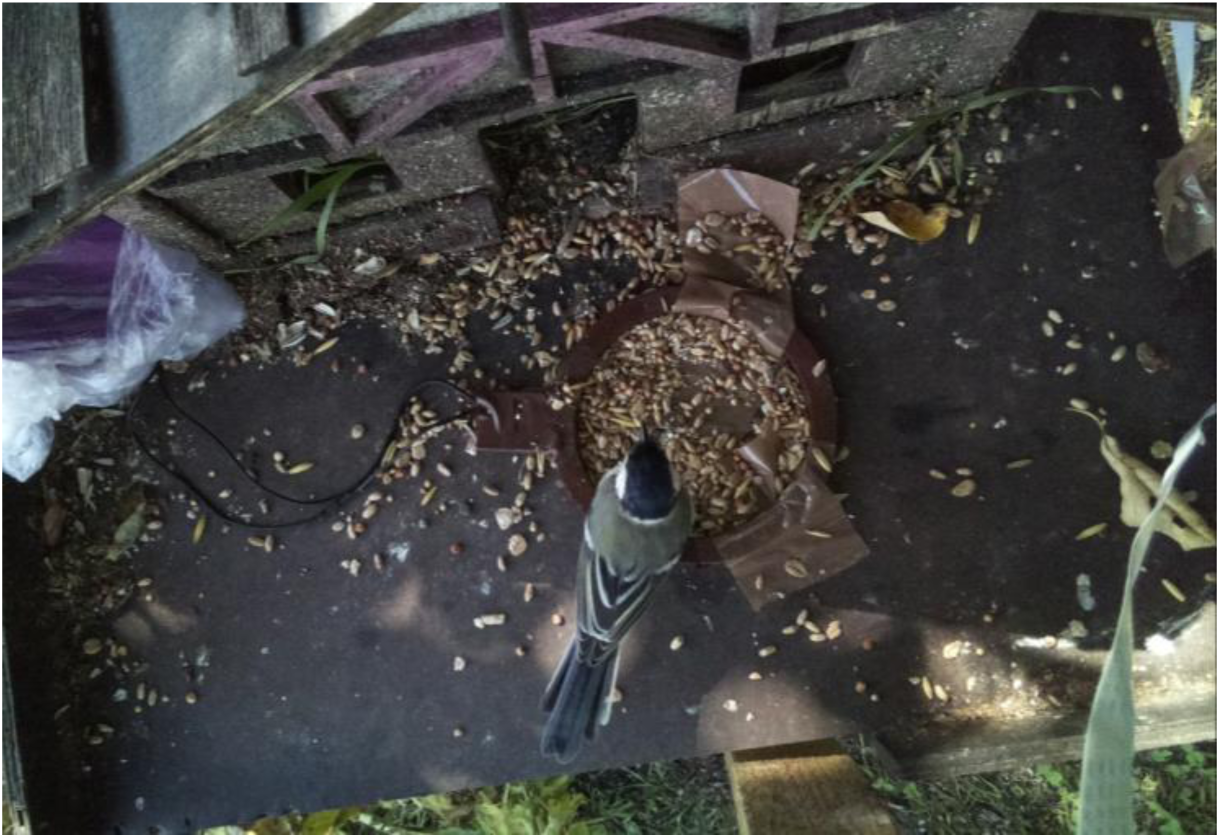
Great tit recorded from a top perspective feeding at a table on top of a RFID antenna.

#### Zebra finches

For the zebra finches we did not have a second setup that differed from the one used to collect the pictures to train a CNN and that could be used for testing the CNN generalization. Therefore, we ran an additional trial which consisted of recording the birds together to see how well the model would predict the identity of the birds when they are in small groups, interacting with each other (Fig. 6). Since these birds did not have any visual tags and it was not possible to distinguish them when in group, we used one flock of three birds and another flock of two birds for each sex to estimate the model’s accuracy by calculating the number of times that the CNN wrongly attributed the identity of a bird as being an individual that is not effectively present in that flock. In order to avoid near-identical pictures, the same procedure as for the validation dataset to select 160 pictures from each trial was used.

**Figure 6.**
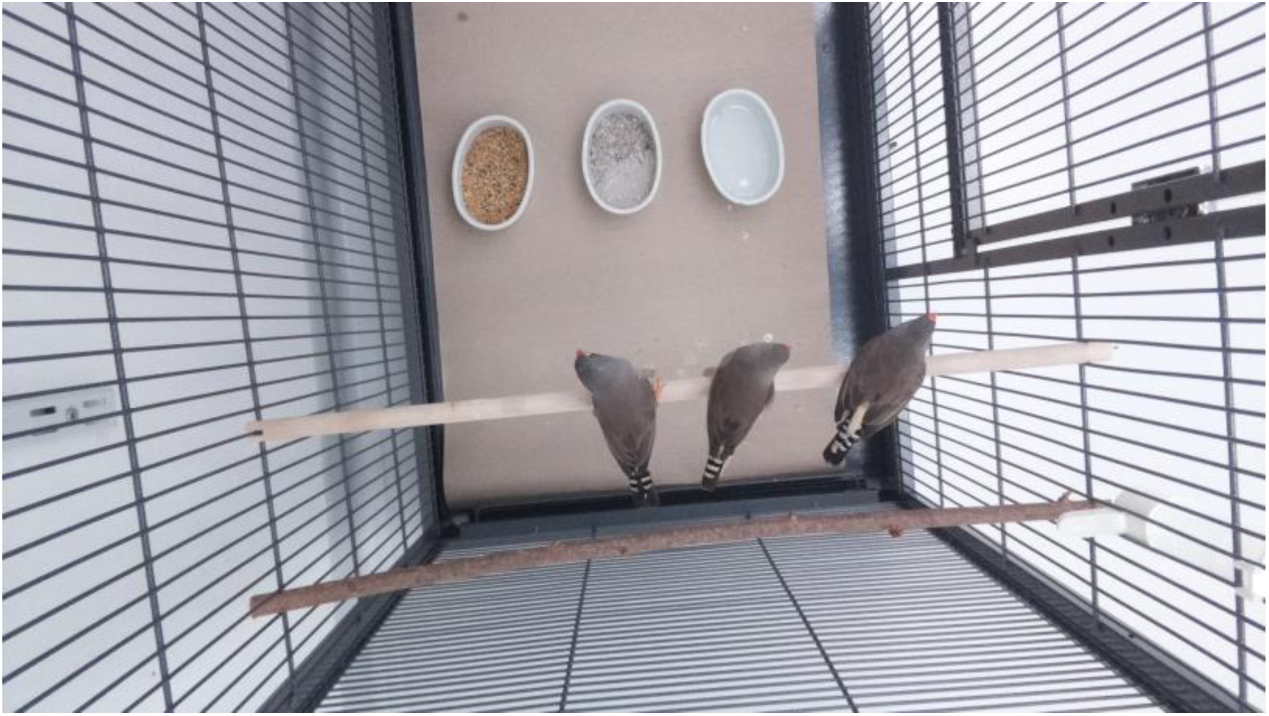
Example of a picture used for the zebra finches’ testing dataset.

### New birds

In wild populations, new individuals can join the population during the course of a study. These new individuals may challenge the performance of a CNN because the model outputs a vector from a softmax layer that indicates probabilities of presence for every individual present during training and the sum of these probabilities is one (see “classification” stage in Fig. 2). In order to study this potential issue we used the already trained CNNs to predict the identity of birds that were not in the training dataset. For the sociable weavers, a scenario in which a CNN was trained to identify a relatively large number of individuals (30) was used to expose the obtained CNN to a small number of new individuals (5). For the great tits the opposite scenario was tested by using a CNN that was trained for a small group of individuals (10) and is exposed to a large number of new individuals (67). For the sociable weavers, we selected 50 pictures of each of the five birds (a total of 250) that were not in the training dataset and 250 random pictures from the pool of birds used during training. For the great tits 250 random pictures were selected from the pool of 67 individuals that were not in the training dataset. We limited the number of pictures from the same individual to a maximum of eight (3.91± 1.67 mean±SD) in order to keep a large number of different individuals in this dataset (64 out of the 67 were used) and randomly selected 250 pictures from the 10 individuals for which the CNN was trained. Shannon’s entropy of each of the distributions was calculated from the classification (softmax) output to empirically determine a confidence threshold to consider a bird as part of the training dataset.

## RESULTS

### CNN

#### Sociable weavers

The model was able to achieve an accuracy of 92.4% (Table 1) after training for 21 epochs. When the model was used to predict the identity in four other contexts, it appears that the accuracy of top perspective’s context was lower (67.5%). After adding blur and noise to the training images, the model achieved a validation accuracy of 90.3%, while successfully increasing the accuracy from the top perspective to 91.6% (Table 1).

**Table 1.** Rate of positive identification when testing in all contexts for the sociable weavers. Right column gives the identification success rate when noise and blurs were artificially added to training images to match the quality of testing images.

| Perspective | Positive identification | Positive identification<br>after adding blur and<br>noise |
| --- | --- | --- |
| Validation | 0.924 | 0.903 |
| Close | 0.926 | 0.926 |
| Far | 0.958 | 0.972 |
| Ground | 0.867 | 0.944 |
| Top | 0.675 | 0.916 |

#### Great tits

The model reached 90.0% accuracy after training for 32 epochs. When using the pictures from the top perspective recording the birds on the table the model correctly predicted the identity of the birds in 85.1% of the pictures.

#### Zebra finches

The model reached 87.0% accuracy after training for 11 epochs with similar accuracies for males and females (85% for males, 88.9% for females). When using the trained model to predict the identity of the birds when they were in small groups the model correctly predicted the identity of a bird present in that group in 93.6% of the time.

### New birds

The entropy of the softmax outputs (i.e. probabilities) was smaller when predicting the identity of birds present in the training dataset, compared to when predicting the identity of new birds (Fig. 7). This is due to the fact that when predicting the identity of a bird from the training dataset, there is usually one that stands out with very high probability (indicating the bird’s identity) and the remaining probabilities are very low (other birds’ identities). In contrast, when predicting the identity of a new bird, the probabilities were usually more equally distributed across all classes, all with low values.

**Figure 7.**
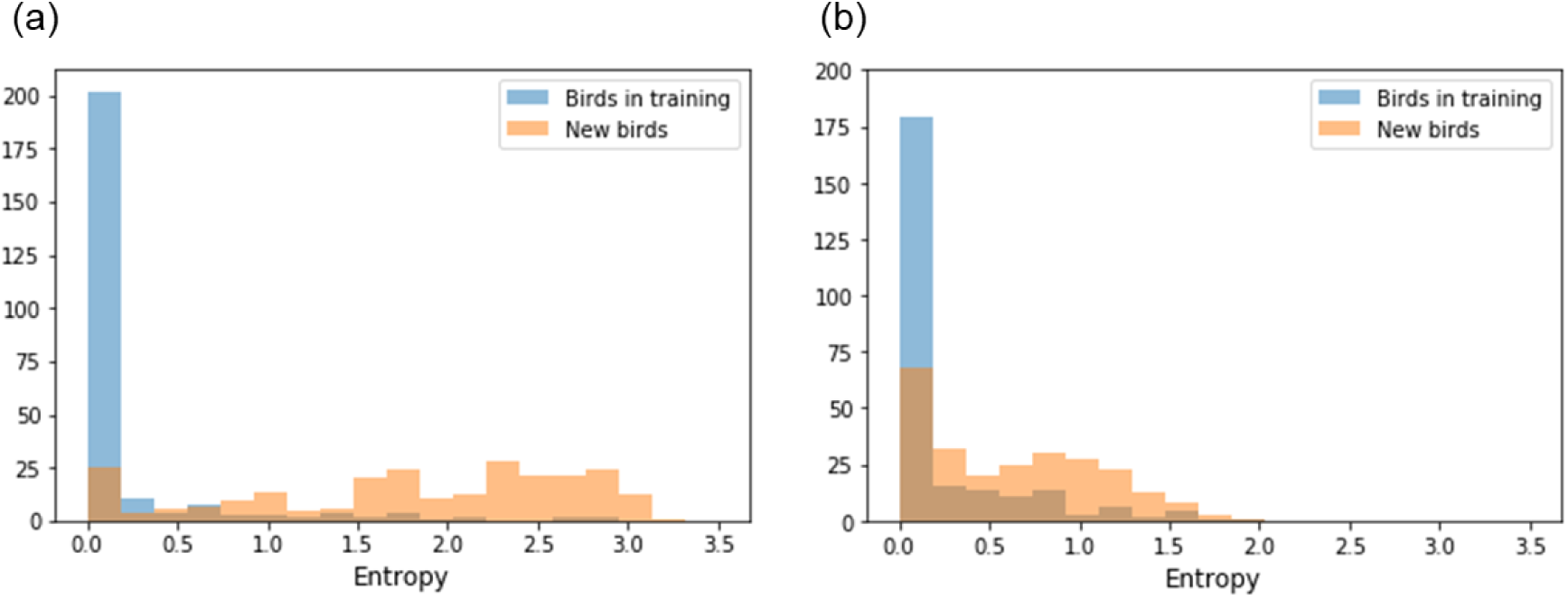
Distribution of the entropies of softmax probabilities when predicting the identity of birds from the training dataset or of new birds for a) sociable weavers and b) great tits.

For the sociable weavers 90% of entropies are below 0.75 when predicting the identity of birds from the training dataset and only 17% of them are under this value when predicting the identity of new birds. This means that with this 0.75 threshold there is a 17% chance that a new bird will be erroneously classified as one of the birds of the training dataset. A value of 17% should be acceptable if new individuals are not common.

For the great tits scenario, in which the appearance of new birds is frequent, defining a simple threshold would not be enough as there is a too much overlap between the birds in the training and the new birds’ entropy.

## DISCUSSION

Deep learning has the potential to revolutionize the way in which researchers identify individuals. Here, we propose a practical way of collecting large labelled datasets, which is currently identified as the main bottleneck preventing the application of deep learning for individual identification in animals (Schneider, Taylor, Linquist & Kremer, 2018). We also demonstrate the steps required to train a classifier for individual identification. To our knowledge, this is the first successful attempt of performing such an individual recognition in small birds. Using data collected with automatized procedures, CNNs proved to be effective for identifying individuals in three different bird species, including two species that are among the most commonly used models in the field of behavioural ecology, and therefore such results highlight the potential of applying CNN to a vast range of research projects. Furthermore, we found high generalization capacities of the trained CNNs, meaning that the rate of successful identification remained high in various contexts. This is particularly relevant as researchers often need to collect data in contexts that may be challenging, from parental behaviour at the nest to dominance interactions at artificial feeders. However, we also show that the models’ performance can become lower when new individuals join the population, especially when new individuals are common..

The first critical step when attempting to implement deep learning is to guarantee that enough training data can be collected to train a model. In this study, for the two wild populations, we showed that we can rely on RFID technology to gather large amounts of automatically labelled data. Since this technology has been increasingly used on birds, we believe that the proposed method for automatizing data collection for deep learning applications could be easily and rapidly implemented in a large number of research programs. Furthermore, the method could be easily extended to other animals and other identification techniques. The main idea is to develop a framework in which the same individuals can be repeatedly photographed, while those pictures are automatically labelled. For example, GPS (e.g., Weerd et al., 2015) or proximity tags technology (e.g., Levin, Zonana, Burt & Safran, 2015) could also be used in combination with camera traps to collect training data. Even with non-electronic tags, it should be possible to design setups to photograph animals automatically, such as by isolating the animals as we showed here with the zebra finches. With the popularization of imaging and sensor technologies, we believe that efficiently collecting a large amount of data should no longer represent a bottleneck preventing the application of deep learning methods such as CNN.

Variation in the recording conditions, for example due to light intensity, shadow or characteristics inherent to the recording quality, should also be taken under consideration as it could limit the model generalization and application ability. Photographing the animals across different times of the day and in different days provides the CNN with a very diverse training dataset making the CNN invariant to such variations. Furthermore, we show here that if the conditions for training are slightly different from the recording conditions in which the CNN is going to be applied, it is possible to artificially modify the pictures used for training in order to simulate the conditions under which the pictures of the context of interest will be taken. Specifically, we used blur and noise transformations in the sociable weaver dataset to improve the generalization capability of our model as the testing images had a lower quality. This confirms that using artificially degraded training pictures can be used to improve CNN generalization capability (e.g. Vasiljevic, Chakrabarti & Shakhnarovich, 2016). Other transformations could potentially be applied on the training dataset. Such transformations should consider the type of images on which the model will be used. For example, if illumination conditions of the training pictures are different from the context of interest, brightness and contrasts transformations could be applied to the training data in order to make the CNN light invariant. This generalization capability is an important novelty of this study compared to previous work on small-animal tracking using computer vision, which have been restricted to standardized conditions and to a fixed number of individuals determined beforehand (e.g. Pérez-Escudero et al., 2014), which are not feasible when working with wild animal populations.

For research questions that do not need long time windows of data collection or that are conducted on species that maintain their appearance with great consistency, collecting training data within a short-period of time might be enough for developing an algorithm for individual identification. However, for longer-term studies and when working with species that have the potential to change their appearance (e.g. moulting in birds), this constitutes a potentially serious limitation. The problem of long-term application of neural network algorithms has been studied in the context of place recognition (e.g. streets recognitions; Gomez-Ojeda et al., 2015); however, to our knowledge, there is still no study addressing the impact of changes in appearance in animals in deep learning-based identification. Currently, we do not know if using training data collected during long periods of time or targeting specific parts (e.g. excluding the wing feathers and considering only the top part of the back, or other body parts such as the flank or the bib) of the birds would make the CNN appearance-invariant by learning more conservative features of the birds that are kept across moulting events. In order to fully address the problem and the potential solutions, pictures of birds collected over longer periods of time and from multiple body parts are needed. However, while these datasets are not available, the automatization of training data collection is an immediate and effective solution, i.e. it is possible to continuously collect training pictures and routinely re-train the CNNs using the new updated dataset.

The arrival of new individuals to the study population is another challenge that needs to be carefully addressed. If these new birds are marked with a pit-tag, the CNN could be updated similarly to the problem of changes in appearance discussed above. If the new individuals are not marked and cannot be captured the problem fits in the anomaly (Chandola, Banerjee & Kumar, 2009) and novelty (Pimentel, Clifton, Clifton & Tarassenko, 2014) detection domain. Here we used a simple approach based on the entropy of classification probabilities, which appeared useful if the CNN was trained on a relatively large number of individuals and if immigrants are uncommon in the population, like in the sociable weaver example. Moreover, the error rate might be reduced if the identification is based on a collection of frames (e.g. pictures extracted from a short video recording of the animal) instead of single picture. However, for some studies, such conditions might not be met and, as we showed for the great tit scenario, where we had a low number of individuals in the training dataset and observed a large number of new birds, other approaches have to be explored when sufficient individual data is available (for example by using Siamese neural networks; Varior, Haloi & Wang, 2016). The field of deep learning progresses due to the existence of large and freely availed databases which are used to try different approaches for a wide range of classification problems. For example, the ImageNet database (Deng et al., 2009) has been used numerous times to create algorithms for object recognition. The LFW dataset (Huang, Mattar, Berg & Learned-Miller, 2008) contains thousands of pictures of human faces to development algorithms for human face recognition and identification. The nordland dataset (Sünderhauf, Neubert & Protzel, 2013) contains footage of more than 700km of northern Norway railroad recorded in different seasons (summer, winter, spring and fall) and has been used to address the problem of place recognition under severe environmental changes. Similarly, biologists aiming at taking advantage of the potential of deep learning need large datasets with labelled pictures of several individuals, taken across different contexts and across different life stages, in order to develop reliable algorithms that are able to cope with the challenges presented here, among others.

Having large datasets will also allow optimizing the CNN performances. Other network architectures (e.g. ResNet; He, Zhang, Ren & Sun, 2016) and different hyper-parameters settings (e.g. learning rate) than the ones used here can yield different, and potentially improved, results. There are also other pre-processing steps that can greatly improve the model training and reduce the number of images needed such as, image alignment (e.g. Deb et al., 2018; Lopes, de Aguiar, De Souza, & Oliveira-Santos, 2017), which could be used to decrease variation in the birds’ pose. Training a CNN encompasses a great deal of trial and error and different systems will present different challenges. Nonetheless, we hope that our work will motivate other researchers to start exploring the possibility of using deep learning for individual identification in their model species, and conduct further work on addressing the constraints of working with birds both in the wild and in captivity (namely moulting and introduction of new individuals). The ability to move beyond visual marks and manual video coding will revolutionise many of the questions we can address by making data collection more efficient, cheaper and faster.

## ACKNOWLEDGEMENTS

Data collection on the sociable weaver population would have not been possible without the contribution of several people working in the field, in particular those who contributed to operate the RFID stations and conduct the annual sociable weavers captures: António Vieira, Rita Fortuna, Pietro D’Amelio, Cecile Vansteenberghe, Franck Theron, Annick Lucas, Sam Perret, and several other volunteers. De Beers Consolidated Mines gave us permission to work at Benfontein Reserve. We also thank Gustavo Alarcón-Nieto and Adriana Maldonado-Chaparro for the assistance with the material needed to collect pictures of the great tits and the zebra finches. Data collection for the sociable weaver data was supported by funding from the FitzPatrick Institute of African Ornithology (DST-NRF Centre of Excellence) at the University of Cape Town (South Africa), FCT (Portugal) through grants IF/01411/2014/CP1256/CT0007 and PTDC/BIA-EVF/5249/2014 to RC and the French ANR (Projects ANR-15-CE32-0012-02 and ANR 19-CE02-0014-02) to CD. This work was conducted under the CNRS-CIBIO Laboratoire International Associé (LIA). ACF was funded by FCT SFRH/BD/122106/2016. FR was funded by national funds through FCT – Fundação para a Ciência e a Tecnologia, I.P., under the Scientific Employment Stimulus - Individual Call - CEECIND/01970/2017. This work benefited from a grant by the Deutsche Forschungsgemeinschaft (DFG grant FA 1402/4-1) awarded to DRF. DRF and HBB received additional funding by the Max Planck Society and the DFG Centre of Excellence 2117 “Centre for the Advanced Study of Collective Behaviour” (ID: 422037984).

## AUTHORS’ CONTRIBUTIONS

ACF, LRS, CD and JPR had the idea of applying deep learning for individual identification in the sociable weaver population and DRF had the idea of applying it to the zebra finch and great tit populations. ACF and LRS developed the RFID and Raspberry Pi based method for automated training data collection. LRS analysed the sociable weaver videos for testing the model generalization capability. RC and CD provided all the required funding, material and access to the individually marked sociable weaver population and DRF to the great tit and zebra finch populations. ACF, HBB and DRF developed the setup to collect pictures of the zebra finches. ACF, HBB collected the data of the zebra finches. ACF collected the data for the sociable weaver and the great tit populations. ACF led the statistical analysis and data pre-processing assisted by FR and JPR. ACF wrote the first draft of the manuscript. All authors contributed to editing and revising the final manuscript.

## DATA ACCESSIBILITY

All scripts and data for reproducing the entire contents of this article are available at https://github.com/AndreCFerreira/Bird_individualID.

